## Supplementary figures and images for "Emergence of Non-Canonical Parvalbumin-Containing Interneurons in Hippocampus of a Murine Model of Type I Lissencephaly"

### Figure S1

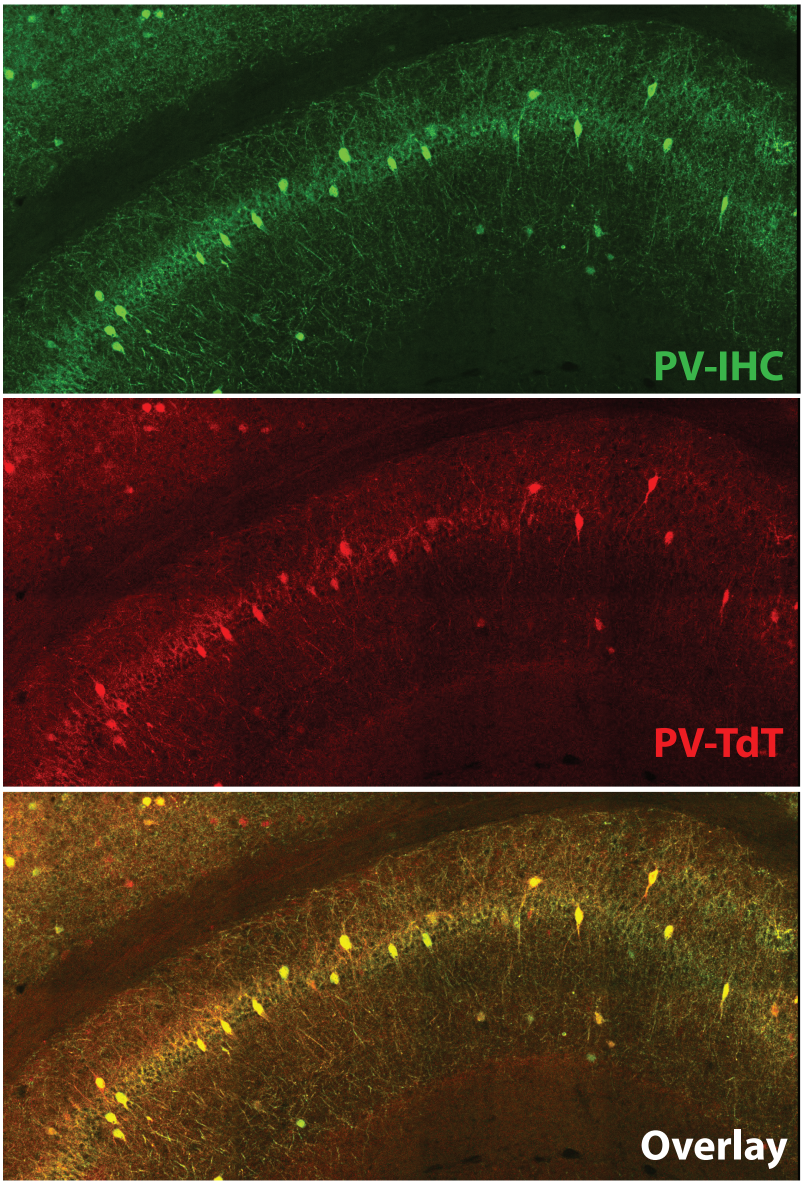

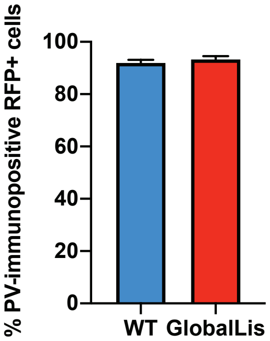

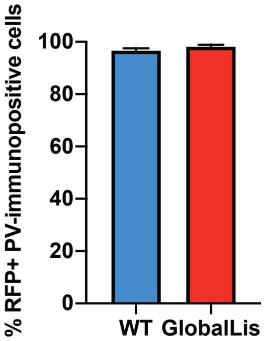

### Figure S2

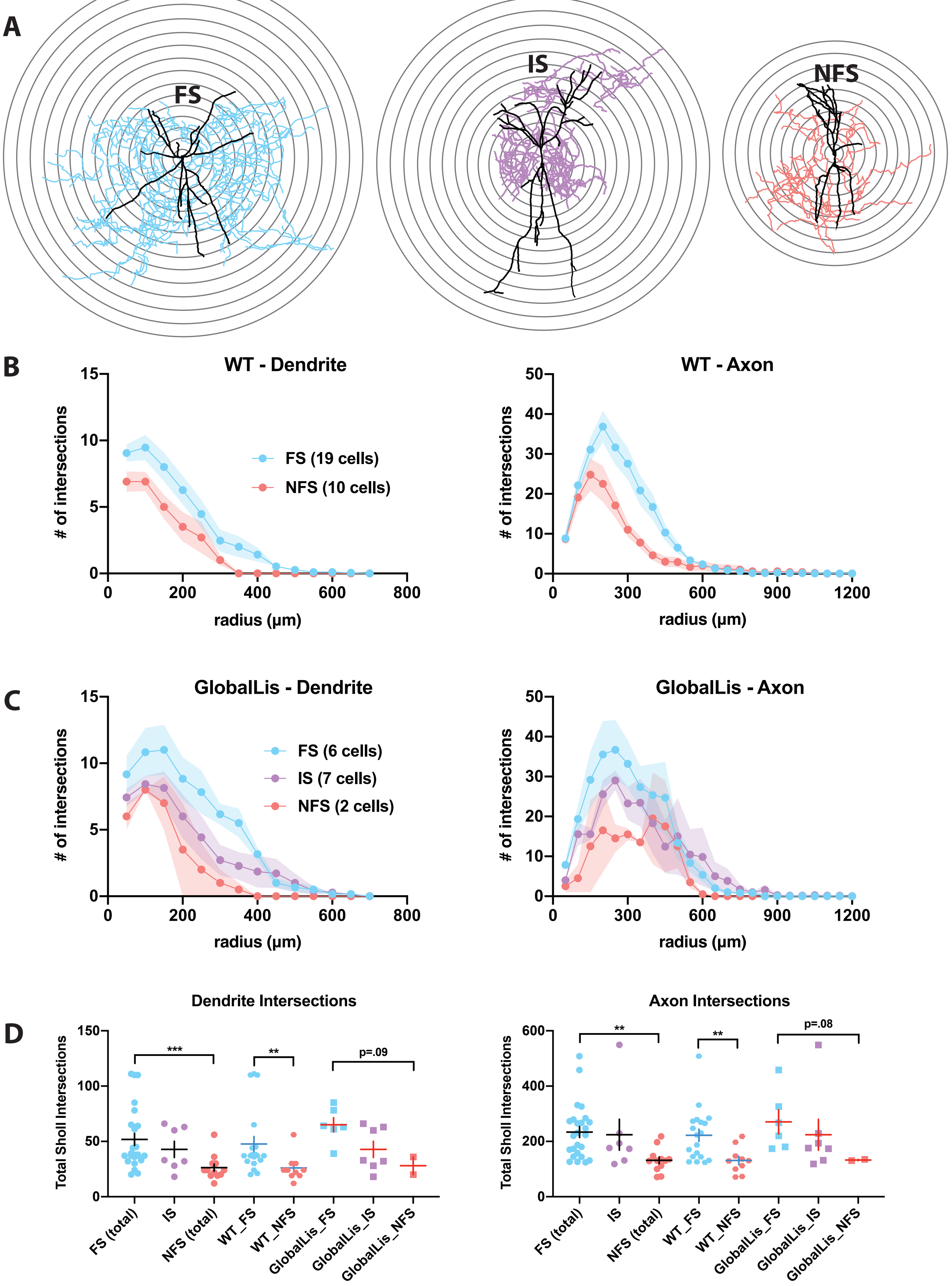

### Figure S3

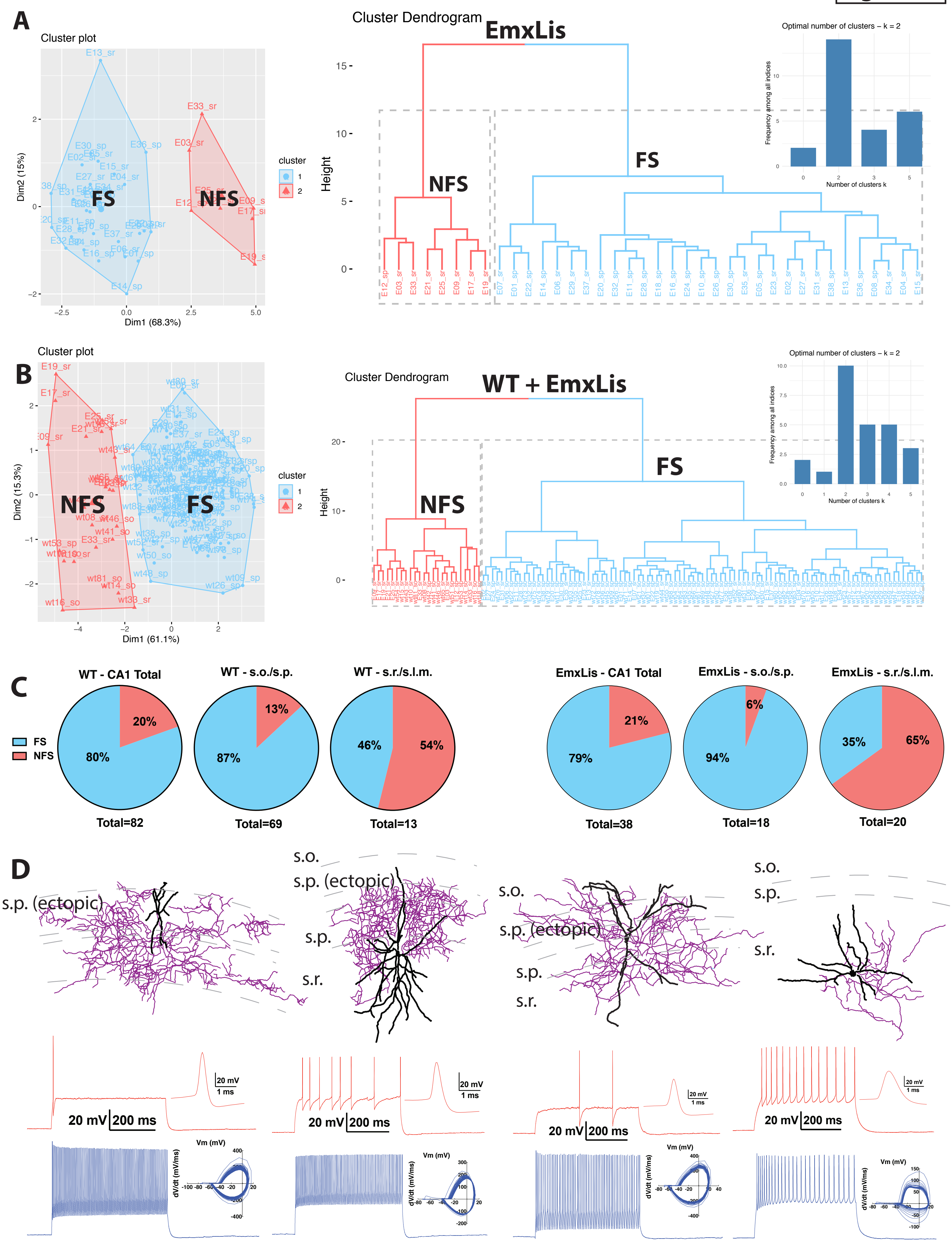

### Figure S4

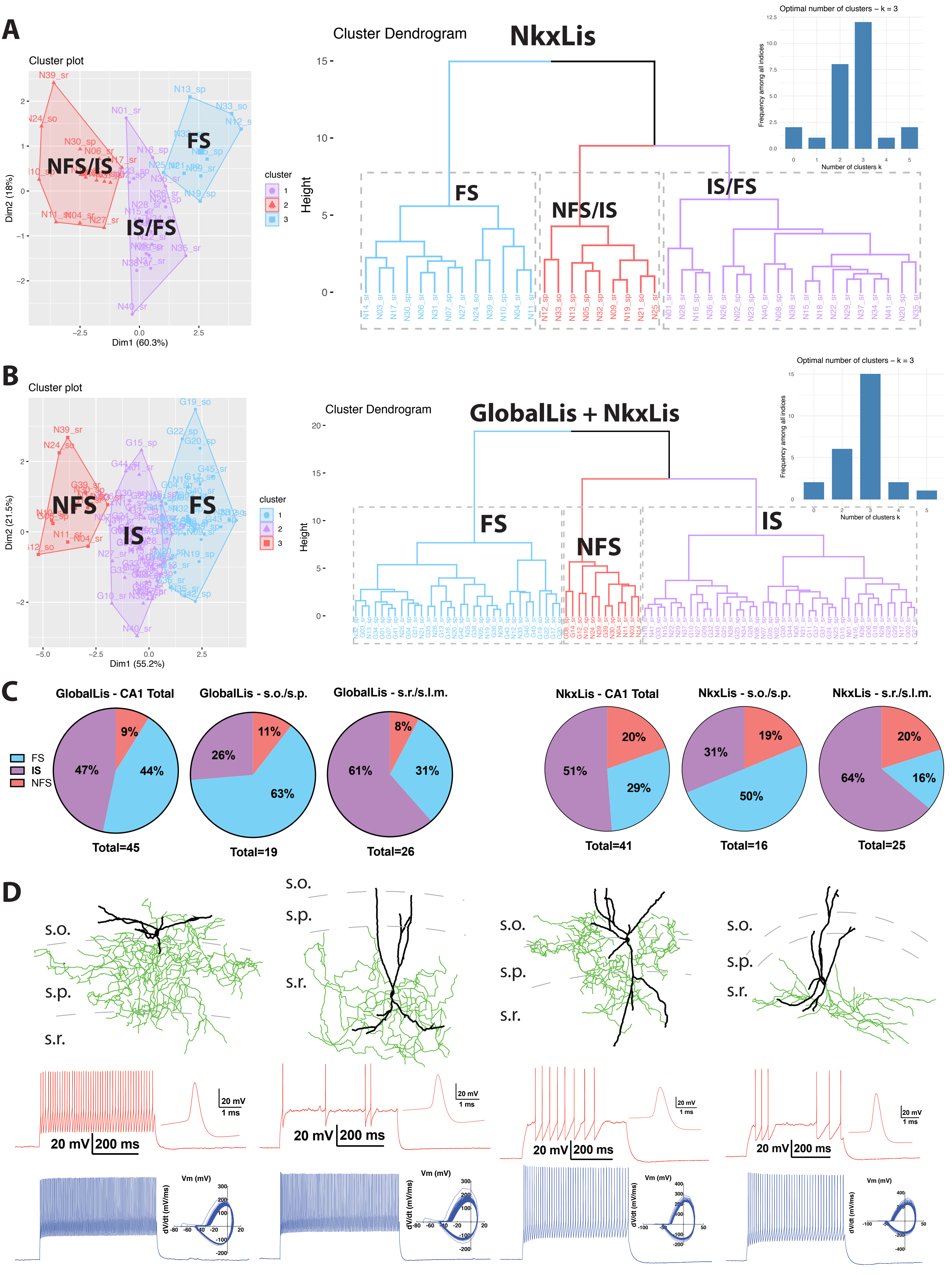

### Figure S5

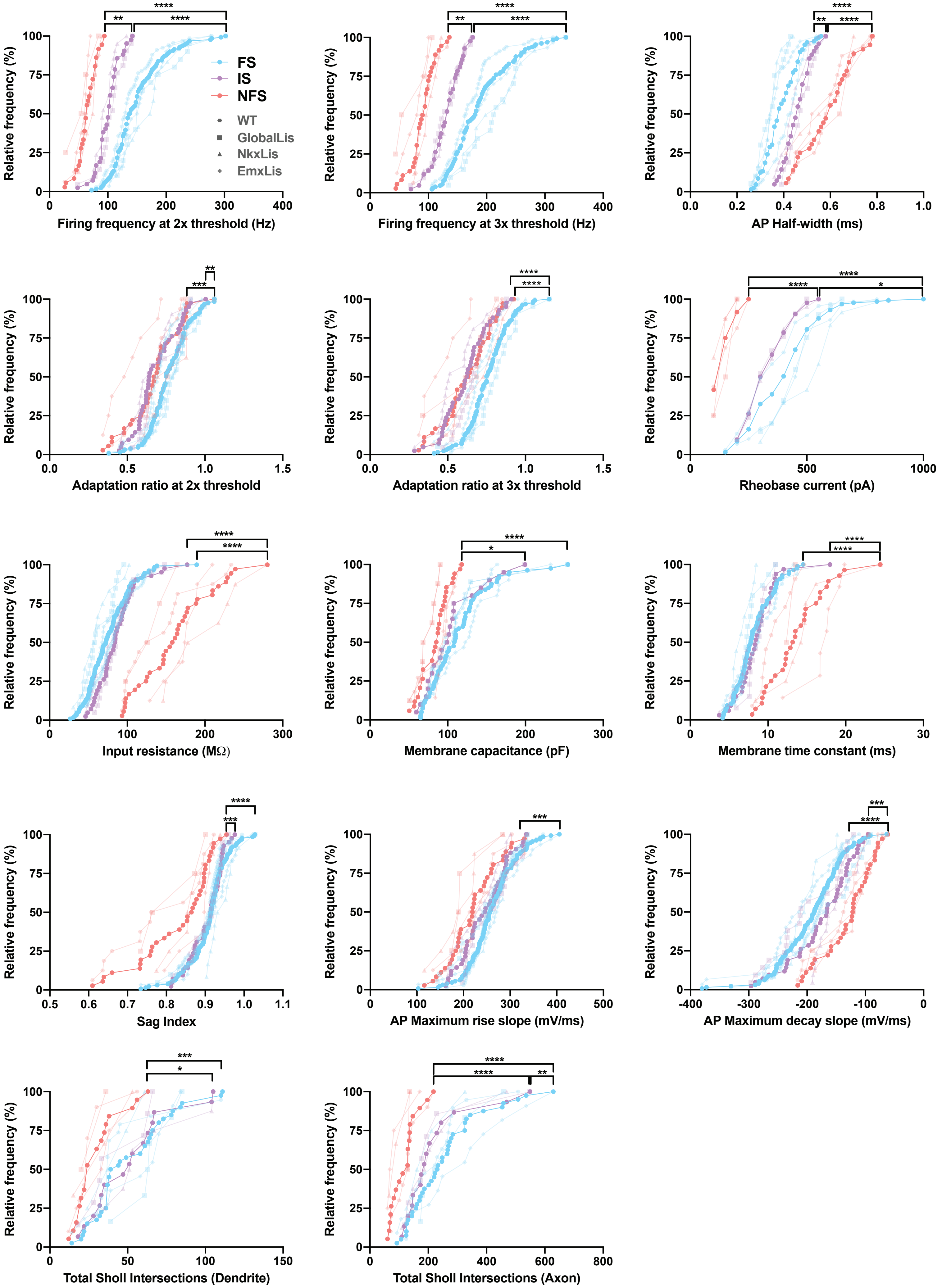

### Figure S6

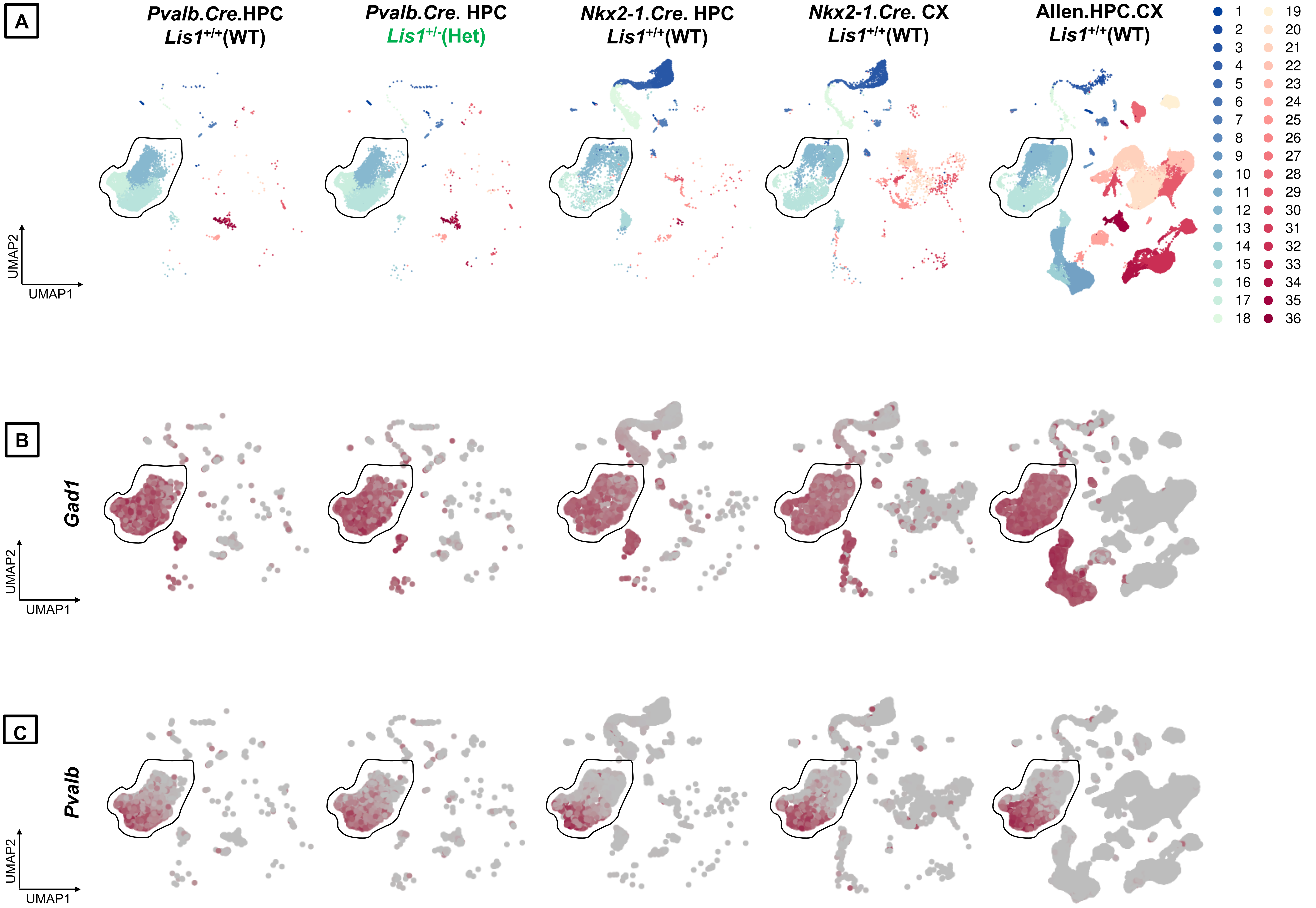

### Figure S7

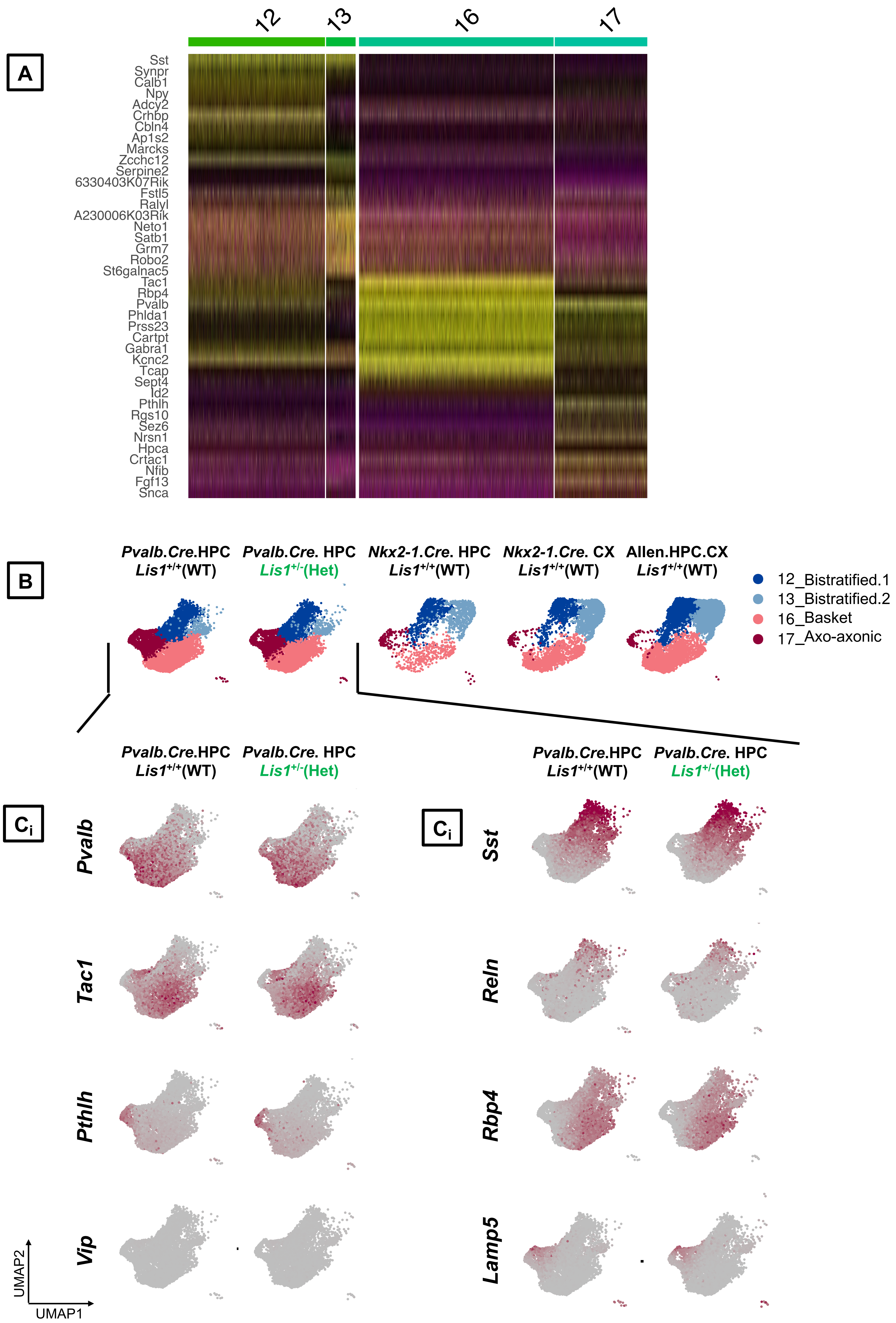
